# Adaptive generalization in pollinators: hawkmoths increase fitness to long-tubed flowers, but secondary pollinators remain important

**DOI:** 10.1101/2024.03.08.584170

**Authors:** Katherine E. Wenzell, Johnathan Y. Zhang, Krissa A. Skogen, Jeremie B. Fant

## Abstract

Selection on floral traits by animal pollinators is important in the evolution of flowering plants, yet whether floral divergence requires specialized pollination remains uncertain. Longer floral tubes, a trait associated with long-tongued pollinators, can also exclude other pollinators from accessing rewards, a potential mechanism for specialization. Across most of its range, *Castilleja sessiliflora* displays much longer corollas than most *Castilleja* species, though tube length varies geographically and correlates partially with hawkmoth visitation. To assess if long corolla tubes reflect adaptation to hawkmoth pollinators, we performed a day/night pollinator exclusion experiment in nine natural populations that varied in corolla length across the range of *C. sessiliflora* and short-tubed members of the parapatric *C. purpurea* complex. We compared the fitness contributions of nocturnal and diurnal visitors, revealing that long-tubed populations visited predominantly by hawkmoths experienced greater fruit set at night, in contrast with short-tubed populations or those visited mainly by diurnal pollinators. Next, leveraging a range-wide multi-year dataset of pollinator visitation to these species, we identify that hawkmoth visitation is associated with increased fitness in long-tubed populations overall, and that long tubes are associated with less diverse visitor assemblages. Thus, long corollas represent an adaptation to hawkmoth pollination at the exclusion of diverse pollinators. Nonetheless, while hawkmoths were scarce in the northern range, secondary diurnal pollinators contributed to fruit set across the range, providing reproductive assurance despite possible trait mismatch. This study illustrates adaptive generalization in pollination modes and that floral divergence may proceed along a continuum of generalized and specialized pollinator interactions.

## Introduction

The diversity of angiosperms, the most species-rich lineage of plants, is frequently linked to their association with animal pollinators (Crepet et al., 2004; van der Niet and Johnson, 2012; Wei et al., 2021). Pollinators drive the diversification of angiosperms by selecting for differing floral traits, which may then act as isolating barriers among plant species, spurring divergence and speciation (Fenster et al., 2004; Johnson, 2006). However, open questions remain regarding the process of pollinator-mediated plant speciation, including how floral traits respond to selection from geographically variable pollinator assemblages, or from distinct pollinator groups within a generalist assemblage of visitors (Kay and Sargent, 2009). Classic models suggest that plants visited by a broad generalist array of pollinators are unlikely to adapt to any one of these pollinators (Waser et al., 1996), and are therefore unlikely to exhibit floral divergence, while other models posit such divergence is possible if the plant adapts to one most effective pollinator (Stebbins, 1970), presumably at the exclusion of other inferior pollinators. Nonetheless, recent work suggests that divergence with generalization is possible via “adaptive generalization” whereby multiple less effective pollinators may still visit and pollinate flowers otherwise adapted to the most effective pollinator, as long as the inferior pollinators do not result in a net fitness cost to the plant (Ohashi et al., 2021). Thus, the evolution of “filter traits,” or those that function to exclude certain less effective pollinator groups provides a framework to investigate the ecological circumstances under which generalist pollinator assemblages may be favored, or when fitness trade-offs may result in specialization (Sargent and Otto, 2006; Miller et al., 2014). Tests of this concept should disentangle how different members of a plant’s pollinator assemblage contribute to its fitness in light of variable floral morphology.

When considered in the context of geographic variation, trade-offs following specialization may result in trait mismatch (Thompson, 2005), wherein traits are maladapted to local conditions, particularly at range edges due to low population density and gene flow (Kay and Sargent, 2009). If fitness trade-offs favor specialization toward the most effective pollinator through the evolution of filter traits, this risks trait mismatch and a lack of reproductive assurance in regions where the most effective pollinator is rare or absent, representing a fundamental risk of specialized pollination systems (Waser et al., 1996). Floral trait shifts that are strongly directional or irreversible (i.e., unlikely to revert to the ancestral state) are expected to more likely result in trait mismatch, wherein conflicting selection pressures may be insufficient to overcome evolutionary constraints (Barrett, 2013). Examples of floral traits expected to be irreversible include loss of function mutations such as loss of floral pigmentation (Rausher, 2008; Ho and Smith, 2016), as well as elongation of nectar spurs or corolla tubes ((Whittall and Hodges, 2007; Barrett, 2013), though see (Wang et al., 2023)). Loss of floral pigmentation and elongated floral tubes are both classically associated with pollination by long-tongued hawkmoths (Faegri and van der Pijl, 1971; Fenster et al., 2004), likely contributing to the characterization of hawkmoth pollination by some authors as an evolutionary dead-end (Tripp and Manos, 2008), despite the tendency of hawkmoth abundance to vary in space and time (Miller, 1981; Campbell et al., 1997). With the directionality of these traits in mind, studies of the pollination ecology of a plant species exhibiting natural variation in pigmentation and floral tube length provide an interesting opportunity to investigate relationships between floral traits and hawkmoth pollination, as well as how such interactions might vary across a species range.

One example of such a species is the widespread and phenotypically variable *Castilleja sessiliflora*. This species occurs throughout the plains of central North America and displays variation in inflorescence color, from pale greenish white in the northern range extent to pale pink in the southern range, with some southern populations bearing either bright pink or yellow flowers (Wenzell et al., 2021). These distinctly colored southern populations are also notable for their shorter corolla tubes, in contrast to the long, protruding corollas that characterize *C. sessiliflora* across much of its range. Parapatric to *C. sessiliflora* at its southeastern range extent in central Texas and Oklahoma is the *C. purpurea* species complex (Nesom and Egger, 2014), which is expected to be closely related to *C. sessiliflora* based on morphological and geographic proximity (Chuang and Heckard, 1991; Tank et al., 2009). Species of the *C. purpurea* complex include the purple-bracted *C. purpurea*, the yellow-bracted *C. citrina*, and the red-bracted *C. lindheimeri* (Nesom and Egger, 2014).

Previous work on variation in floral traits and floral visitors across the ranges of *C. sessiliflora* and the *C. purpurea* complex revealed that, despite being visited by broad assemblages of generalist pollinator groups, divergence in several key floral traits mirrored differences in visitation from local pollinators (Wenzell et al., 2023b). Several of these floral traits were related to pollinator attraction, such as color (butterflies were associated with purple and pink flowers, while bumblebees were more likely to visit pale, yellow flowers and avoid red), while other traits were related to mechanical fit and efficiency of pollen transfer (e.g., small bee visitation was associated with less exserted stigmas, expected to increase pollen transfer).

Interestingly, hawkmoths visited all four species at comparable rates on average and were not associated with longer corollas, as had been expected because *C. sessiliflora*’s floral traits align with classical conceptions of moth-pollinated flowers (long, narrow floral tubes and pale pigmentation (Faegri and van der Pijl, 1971)), and because hawkmoths were the predominant visitor to populations of *C. sessiliflora* in the southern half of its range (despite their absence in the north). Nonetheless, Wenzell et al., (2023) assessed only floral visitation, and thus could not directly assess the potential for floral visitors to exert selection on these variable traits, or to disentangle whether visitation and pollination may show different patterns (Fenster et al., 2004). For instance, hawkmoths may show no preference for visiting long-tubed flowers, as their long proboscis can access nectar in both short and long corollas; however, they may prove to be more efficient pollinators to long-tubed flowers, as they may better contact anthers and stigma (Fulton and Hodges, 1999; Whittall and Hodges, 2007; Boberg et al., 2014).

To test the hypothesis that long corolla tubes could increase pollination efficiency by hawkmoths, we performed a pollinator exclusion experiment in nine natural populations that varied in corolla length to assess whether the contribution to fruit set of nocturnal visitors (primarily hawkmoths) was greater than that of diurnal visitors in populations where corollas were long. This day-versus-night pollinator exclusion experiment asks: Q1) Do long floral tubes improve pollination from nocturnal pollinators? We hypothesize they do, based on long corolla tubes both improving contact between hawkmoths’ bodies and the sexual organs of plants (thus increasing pollen transfer and resulting in greater seed set to long-tubed plants) and acting as a filter trait to exclude visitation from pollinator groups that cannot reach nectar rewards at the base of long tubes. If supported, we expect both that long-tubed populations with more frequent visitation from hawkmoths have higher reproductive success, and that long-tubed populations will have a less diverse suite of floral visitors. In this study, we test these latter two expectations by leveraging a range-wide dataset of floral traits and multiyear pollinator visitation data in both *C. sessiliflora* and the shorter-tubed *C. purpurea* species complex (Wenzell et al., 2023) to ask Q2) Do long-tubed populations of *C. sessiliflora* experience higher reproductive success when visited primarily by hawkmoths? And Q3) Are long corolla tubes associated with less diverse assemblages of floral visitors? Thus, this study assesses the relative fitness contributions of different pollinator groups in relation to natural variation in a key floral trait impacting pollinator fit across geography, providing a thorough investigation of how geographic mosaics shape floral divergence along a continuum of pollinator generalization and specialization.

## Methods

### Study system

*Castilleja sessiliflora* (Orobanchaceae) is a perennial, hemiparasitic forb that occurs widely throughout the Great Plains of central North America (Figure 1). Flowers are open day and night for several days while stigmas are receptive, borne on indeterminate inflorescences, and produce nectar rewards at the base of corolla tubes (pers. obs). Flowers consist of a long, protruding corolla (atypical of the genus) and a lobed calyx subtended by a bract which is typically green, though all of these floral tissues can vary in color from white-green (common in the northern portion of the range), to pale pink (common in the southwest), and bright pink and yellow (observed at several populations in the southern range extent, Fig. S1; Wenzell et al., 2021). These distinctly colored bright pink and yellow-flowered populations also bear shorter corollas and are visited by distinct pollinator assemblages compared to other nearby populations of *C. sessiliflora*, potentially representing distinct pollination ecotypes (Wenzell et al., 2023). Across the rest of the range, populations of *C. sessiliflora* bear long corollas (Fig. 1B; typically greater than 40-45mm in length on average) and are visited predominantly by hawkmoths (particularly *Hyles lineata*, Fig. 1C) and small, solitary bees in the southwest; while in the northern range, visitors are mainly small bees with occasional bumblebees.

**Figure 1.**
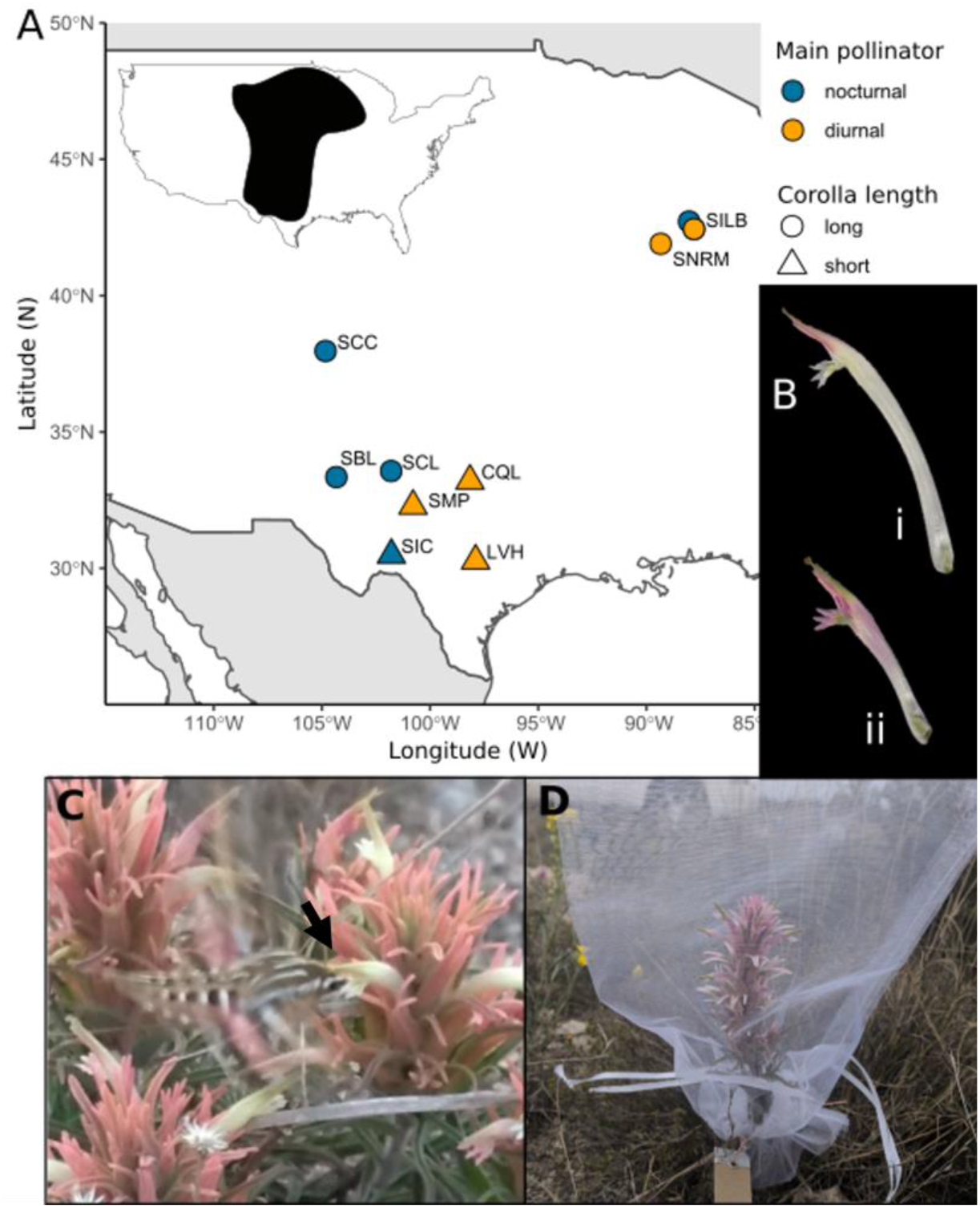
Day/night pollinator exclusion experiment. **A**) Focal populations used in pollinator exclusion experiment; inset shows species distributions. Species identity of populations is denoted by the first letter of the population abbreviation (S= *C. sessiliflora*, C= *C. citrina*, and L= *C. lindheimeri*). Populations represented by circles have long corolla tubes; those shown by triangles have short tubes. Blue points show populations where the most frequent visitor was nocturnal (*Hyles lineata* hawkmoths in all cases); orange points are populations where the most frequent visitor was diurnal. **B**) Representative photos of long (i, SBL) and short (ii, SIC) corollas. **C**) *Hyles lineata* hawkmoth probing a long corolla at SBL. Arrow shows the hawkmoth’s head contacting the plant’s reproductive organs at the mouth of the corolla. **D**) Inflorescence bagged to exclude pollinators at SIC.

To provide a more thorough comparison of short- and long-tubed populations, we chose to include populations of the parapatric *C. purpurea* complex in the pollinator exclusion experiment, due to their geographic proximity, similarity in corolla length to short-tubed populations of *C. sessiliflora* (Fig. 2A), and largely overlapping pollinator assemblages. The *C. purpurea* species complex (comprising *C. purpurea, C. citrina*, and *C. lindheimeri*) occurs in Texas and Oklahoma, USA (Nesom and Egger, 2014) and are also perennials with almost identical floral morphology to that of *C. sessiliflora*, with the exception of shorter corollas and showier, wider floral bracts. Pollinator visitation to these species is largely generalized and includes many of the same major functional groups that visit *C. sessiliflora*: Hymenoptera (bumblebees, other large bees, and small, solitary bees) and Lepidoptera (butterflies and hawkmoths), in addition to hummingbirds, particularly to *C. lindheimeri* (Wenzell et al., 2023). Throughout the paper, the species identity of focal populations is denoted by the first letter of population abbreviations: population codes beginning with “S” are *C. sessiliflora*, “C” denotes *C. itrina*, and “L” denotes *C. lindheimeri*. While a population of *C. purpu ea* was initially included in the exclusion experiment, this population was later excluded due to herbivory (see below).

**Figure 2.**
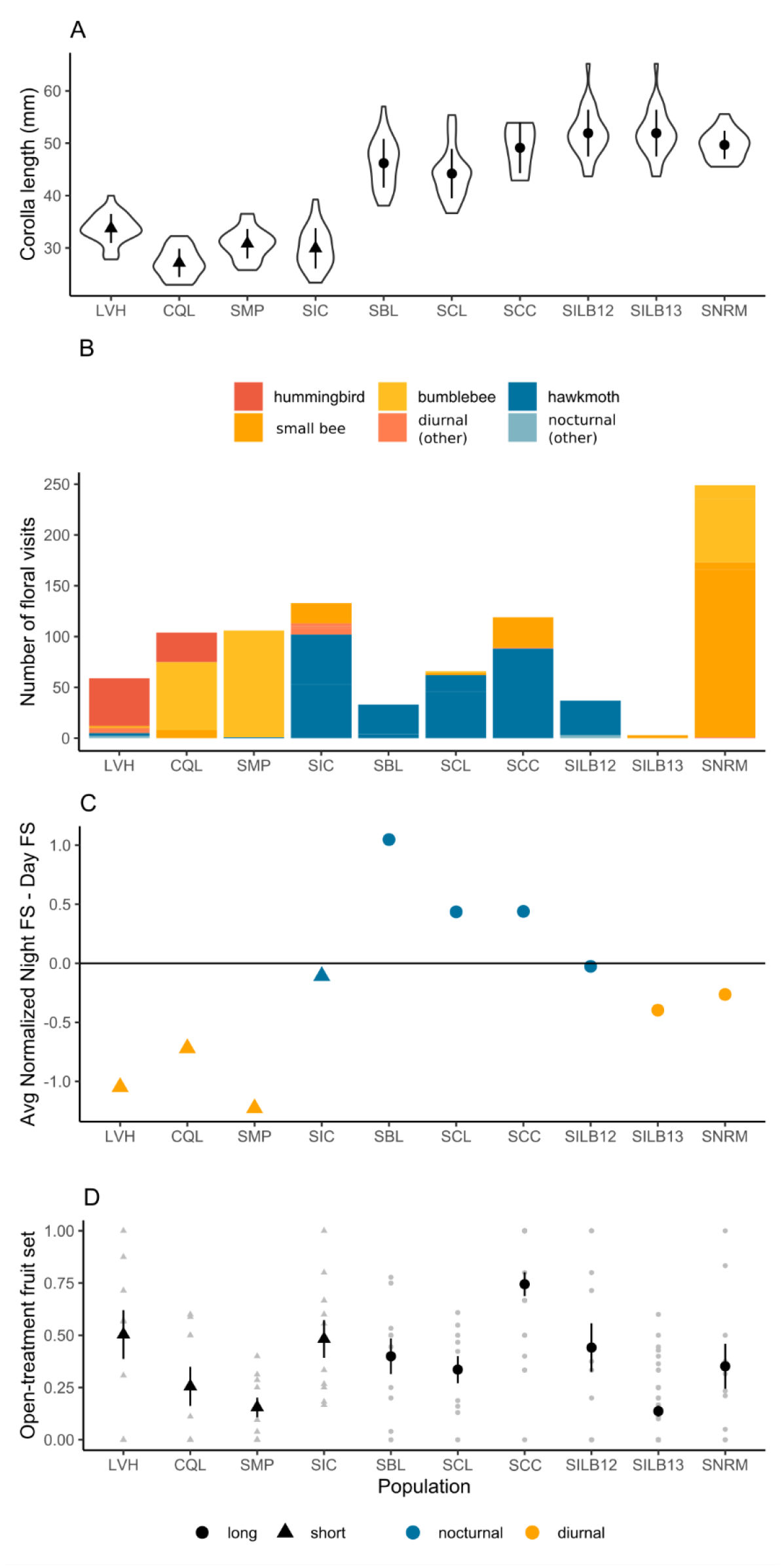
Corolla length, pollinator visitation, and fruit set by population. **A**) Corolla length measurements by population, points show mean +/-SE. Note SILB12 and SILB13 share the same floral measurements taken from population SILB in 2017 (values plotted twice to align with B, C, and D). **B**) Number of floral visits by pollinator functional group observed during exclusion experiments. Diurnal groups are shown in warm colors, nocturnal groups in cool colors. **C**) Difference in average fruit set to night-open and day-open plants [normalized by average fruit set to open-treatment plants: i.e., (Night FS / Open FS) – (Day FS/ Open FS)]. **D)** Fruit set to open-treatment plants: individual values (grey points) and mean +/-SE (black points and bars). Shape of point shows corolla length category and color shows activity period of the most frequent visitor. For all plots, populations are approximately ordered by species and from south to north.

To facilitate pollinator exclusion experiments, the breeding system of *C. sessiliflora* was assessed in several ways. In 2019, flowering *C. sessiliflora* plants growing in a common garden at the Chicago Botanic Garden, Glencoe, IL, were bagged to exclude pollinators to assess autonomous seed production, and a subset of these flowers were hand-pollinated with self-pollen to assess self-fertilization. Fruit set of these flowers was recorded when fruits were mature. Plants were grown from seed directly collected from wild populations across the geographic range of *C. sessiliflora*, including three populations in which pollinator exclusion experiments were performed (SMP, SCC, SILB; see below).

Additionally, in 2012-2013, flowering plants growing in three natural populations in Illinois and Colorado (SILB, SCC, and SDC) were bagged to exclude pollinators, and fruit set was later recorded (see below for additional details).

### Pollinator visitation and floral trait data

For the range-wide, multiyear pollinator visitation dataset previously presented by Wenzell et al. (2023), pollinator observations were conducted at 23 populations across the distributions of *C. sessiliflora* and the *C. purpurea* complex for 1-3 years in March-June 2017-2019 (see Wenzell et al., 2023 for detailed methods). Floral visitors to the focal *Castilleja* sp. were recorded throughout daylight hours and at dusk (until past last light) over one 24-hour period per population per year, and visitors were recorded to functional group (e.g., Fenster et al., 2004) over 440 minutes of observation time per population per year. Floral traits were quantified at these populations from thirty individual plants per population. Floral morphological measurements (including corolla length measurements utilized in this study; Fig. 2A) were taken from two open, receptive flowers per individual, measured from scans taken of fresh tissue. Floral color of inflorescences was assessed *in situ* using Royal Horticultural Society color charts, converted to Red-Green-Blue values (Wenzell et al., 2021, 2023). For additional populations sampled in 2012-2013, pollinator visitation was recorded to focal observation plants following the same method described above, though total observation time varied: 860 minutes of observations were recorded over 2 days for population SCC, 390 minutes for SILB in 2012, and 360 minutes for SILB in 2013 (recorded over one day per year). Additionally, floral traits were measured at SCC in 2016 in the manner described above but for a smaller sample size: two flowers each from four individuals. Given these minor differences in how visitation data and floral trait data were collected in 2012 and 2013, these data were excluded from analyses using the range-wide datasets presented in Wenzell et al., 2023 (described below).

### Do long floral tubes improve pollination from nocturnal pollinators?

We performed a day versus night pollinator exclusion experiment in nine natural populations that varied in population mean corolla length and represented the geographic range of *C. sessiliflora* and members of the *C. purpurea* complex (Fig. 1A). Focal populations were selected to represent the range of variation in corolla tube length in these taxa (Fig. 1B; Wenzell et al., 2021) and included five long-tubed populations of *C. sessiliflora*, from the southern (SBL in eastern New Mexico, SCL in west Texas, and SCC in southeastern Colorado) and northern portions of the range (SNRM and SILB in northern Illinois), and four short-tubed populations: two populations from the southern range extent (the pink-flowered population SIC and the yellow-flowered SMP in southwestern Texas), and one population each of the yellow-flowered *C. citrina* (CQL) and the red-orange *C. lindheimeri* (LVH), both from central Texas (Fig. S1). Initially, an exclusion experiment was also performed at a population of *C. purpurea*, though fruit set could not be accurately recorded due to extremely high levels of herbivory to treatment flowers; thus this site was excluded from analysis. For this experiment, populations with “short” corollas are considered those with a population mean corolla length of less than 34 mm (all *C. purpurea* complex populations and SIC and SMP), and populations with “long” corollas were those with a population mean corolla length value greater than 44 mm (all other *C. sessiliflora* populations), a value which represented a natural break of 10 mm in the data (Fig. 2A; Wenzell et al., 2021). The experiment was conducted during peak flowering period in March - June of 2012 (SCC and SILB12), 2013 (SILB13, the same site sampled in 2012), or 2019 (all 7 remaining populations).

At each population, at least 24-30 flowering plants were bagged with wire cages and mesh (bridal veil) exclusion bags (Fig. 1D) approximately one week prior to the experiment. If plants were bagged less than one week prior to treatments, the most recent open flowers prior to bagging were marked with permanent marker, and flowers up to this point on the inflorescence were excluded from the treatment (since they may have been visited by unknown pollinators prior to bagging). In 2019, treatments were administered by randomly assigning ten plants each to one of three treatment groups: day-open, night-open, and an open-pollinated control. In 2012 and 2013, these same three treatments were applied to one randomly assigned stem each on the same individual plant. Open-pollinated control plants were left unbagged for both day and night for the duration of the experiment (48 hours). Flowers of day-open plants were un-bagged and exposed to pollinators during daylight hours (0.5-1.5 hours post-sunrise, depending on ambient temperature and local pollinator activity, until 0.5 hours before sunset) and were bagged at night to exclude pollinators for two consecutive day/night cycles. Night-open plants were unbagged at night (30 minutes before sunset to 0.5-1.5 hours post-sunrise) and bagged during the day for the same period. At populations sampled in 2012 and 2013 (SCC, SILB12 and SILB13), a fourth treatment was also included, for which plants were bagged and pollinators excluded for the entire treatment period to assess the ability of plants to autonomously reproduce (see above and Supplemental Table S2).

Because *Castilleja* have indeterminate inflorescences, we tracked the flowers included in the treatment as follows: colored wire was tied around each flowering stem below the lowermost open, receptive flower included in the treatment. When treatment plants had multiple stems, each flowering stem was marked with a different color of wire. After the 48-hour treatment period, the number of receptive flowers (from the wire up to the newest receptive flower) were counted and recorded for each stem; plants were then covered with mesh bags to exclude any subsequent pollinators and tagged with an identifying code (Fig. 1D). After 10-14 days (when fruits were maturing and enlarging ovaries were detectable), treatment stems were collected and the number of filled fruits above the wire was counted in the lab. Fitness (i.e., fruit to flower ratio) was calculated as the number of filled fruits above the wire per stem divided by the number of flowers open during the treatment period per stem for each plant. A fruit or ovary was considered filled or enlarged if it contained any seeds or ovules that appeared to be developing into seeds. The proportion of fruit set was used as the response variable in all analyses.

Pollinator observations were conducted at each population (as described above and in Wenzell et al., 2023) during the experimental window, to document the local pool of floral visitors during the time of the experiment. Pollinator functional groups were categorized as either primarily nocturnal (hawkmoths or rarely observed other moths) or diurnal (small bees, bumblebees, hummingbirds, and other uncommon diurnal visitors including bee-flies, butterflies, and hoverflies; Wenzell et al., 2023). The pollinator functional group that contributed the greatest number of floral visits to a population was considered the predominant visitor to that population.

In R (R Development Core Team, 2021), we performed a generalized linear mixed model (GLMM) with package glmmTMB (Magnusson et al., 2020), using data from all experimental populations to test the effect of treatment on proportion fruit set, as well as whether corolla length or identity of the most common floral visitor interacted with treatment to affect fruit set. Proportion fruit set (fitness) was the response variable, and treatment (day-open, night-open, or fully open), corolla length (short or long), and activity period of the most frequent floral visitor to that population during the observation window (diurnal or nocturnal) were included as predictors separately and with an interaction term. Population, individual plant, and year were included as random effects, and the model used a betabinomial distribution weighted by the number of open flowers. The betabinomial error distribution was selected after running the same model with a binomial distribution and comparing the resulting models’ AIC values (the betabinomial distribution resulted in a lower AIC value and was therefore selected). This model was then analyzed using a Type II Wald chi-square test using the Anova() function in R package car (Fox et al., 2020). Model fit and residuals were assessed using package DHARMa (Hartig, 2016), by visualizing scaled residuals based on 1000 simulations, and assessing evidence for zero inflation and significant deviations from the expected distribution based on the fitted model, which were not found.

To directly compare differences in fitness among treatments given contrasting expectations based on corolla length and predominant visitor, we subsetted the full dataset based on corolla length category and major pollinator category, resulting in four subsets: 1) populations with long corollas whose most frequent visitor was nocturnal (i.e., hawkmoths), e.g., “long-nocturnal”, 2) populations with short corollas and predominantly nocturnal visitors (“short-nocturnal”), 3) long-tubed populations visited by diurnal pollinators (“long-diurnal”), and 4) populations with short corollas predominantly visited by diurnal pollinators (“short-diurnal”). We then performed GLMMs on these subsetted datasets to assess if fruit set varied based on pollination treatment (day-open, night-open, or fully open) with population and individual plant included as random effects, followed by Type II Wald Chi-square tests. Pairwise comparisons were then performed using package lsmeans ((Lenth, 2018)) with Tukey adjustments. As described above, model fit and residuals were assessed with package DHARMa, and models were run with both binomial and betabinomial error distributions, and the model with the lowest AIC was selected. This resulted in betabinomial models for long-nocturnal and short-diurnal populations, and binomial models for short-nocturnal and long-diurnal populations.

### Do long-tubed populations experience higher fitness overall when visited by hawkmoths?

To assess if evidence of long corolla tubes and potential hawkmoth-adaptation was generalizable beyond the focal populations of our exclusion experiment, we used a range-wide multiyear dataset (presented and described in Wenzell et al., 2023) to examine whether average fitness varied with respect to corolla length and floral visitors at the population level. We used GLMMs with a betabinomial distribution, weighted by number of recorded flowers, with population as a random effect. The response variable was fitness (proportion fruit set), and the predictor was the identity of the most common floral visitor recorded to that population in the same year in which fruit set was measured, categorized as either hawkmoths (nocturnal) or other diurnal visitors. Populations were grouped into two categories of corolla lengths: short corollas (those with a population mean value of 38mm or less) and long (those with a population mean value of 41mm or greater), and separate models were run on data for long- and short-tubed populations. This resulted in 166 observations from 5 populations (all *C. sessiliflora*) for the long-tubed dataset, and 365 observations from 12 populations (3 populations of *C. sessiliflora* and 9 populations of the *C. purpurea* complex) for the short-tubed dataset. Model fit and residuals were assessed with package DHARMa as described above.

### Do long corolla tubes act as a filter trait to limit the diversity of floral visitors?

To summarize levels of diversity (i.e., generalization) in floral visitors among populations, we calculated the Inverse Simpson’s Diversity Index using number of floral visits by pollinator functional group (adapted from Lázaro et al., 2009). Inverse Simpson’s Diversity Index is less sensitive to rare occurrences than other indices, and thus was chosen to avoid weighting visits from uncommon visitors (Lázaro et al., 2009). Values were log-transformed to approach normality, though data still include a high number of zeroes due to datapoints (i.e., observations in a given population in a given year) in which only one pollinator functional group was recorded. We assessed whether visitor diversity varied with respect to corolla length using a GLMM (with package glmmTMB (Magnusson et al., 2020)) with a Gaussian error distribution and year as a random effect. A fixed effect was included for observation dataset type, based on whether visits were recorded only to designated focal plants or a wide view of the population (see Wenzell et al., 2023 for details). Populations were categorized as either having long (> 41 mm) or short (< 38 mm) corolla tubes on average as described above, and these categories were used as a categorical predictor in the model. Model fit and residuals were checked using package DHARMa (Hartig, 2016), as described above. Based on the large number of meaningful zeroes in the dataset, we chose to use the package glmmTMB, which specifically handles zero-inflated data in GLMMs, and we included a zero inflation parameter of 1 and specified a BFGS optimizer. The model was then analyzed using a Type II Wald chi-square test using the Anova() function in R package car (Fox et al., 2020). Because the model was found to be zero-inflated using the testZeroInflation() function in package DHARMa, in addition to the steps taken above, we also performed a nonparametric Kruskal-Wallis test to test if pollinator diversity value (averaged across dataset within population) varied by categorical corolla length. Finally, given the wide geographic spread of populations and the fact that nearly all populations in the northern portion of the study range have long corollas, we also examined whether geography could explain any difference in diversity of floral visitors by performing another GLMM and Kruskal-Wallis test as described above with population latitude as the predictor, rather than corolla length.

## Results

### Self-incompatibility of *C. sessiliflora*

Pollination treatments to *C. sessiliflora* grown in a common garden revealed that bagged and hand-self-pollinated flowers produced almost no fruits, consistent with a largely self-incompatible breeding system (Supplemental Table S1). Of 33 flowers that were hand-self-pollinated, 0 fruits were produced (0% selfed fruit set), while only 1 fruit was produced out of 105 flowers (<1%) that were bagged and unmanipulated. Similarly, very few fruits were produced by bagged flowers in natural populations: out of a total of 497 receptive flowers across 82 individuals, only 3 fruits were produced (a fruit set rate of 0.6%), all of which occurred in one Colorado population, SCC (Supplemental Table S2).

These findings suggest that *C. sessiliflora* is largely self-incompatible, indicating that fruit set can be attributed to activity by pollinators in our exclusion experiment. Nonetheless, we acknowledge that these experiments did not directly test self-compatibility in *C. citrina* and *C. lindheimeri*, which were used in exclusion experiments and also rely on this assumption.

However, we note that aniline blue staining of self-pollinated pistils of *C. citrina* and *C. lindheimeri* (from natural populations where exclusion experiments were performed) revealed comparable levels of pollen tube germination and subsequent termination in styles as in *C. sessiliflora* (from populations found to be self-incompatible in the common garden experiments), suggesting these species maintain a similar breeding system as their putative close relative *C. sessiliflora* (J. Zhang and K. Wenzell, unpublished data). Further, no evidence of fruit set to flowers produced after the exclusion experiment (when flowers at the top of the inflorescence were completely bagged) was detected in either *C. citrina* or *C. lindheimeri*, providing additional support that observed fruit set can be attributed to pollinators.

### Hawkmoth pollinators confer higher fruit set to long-tubed populations at night

#### Pollinator visitation

Observations of floral visitors during the exclusion experiment revealed a diversity of diurnal pollinator functional groups visiting short-tubed populations in the south (e.g., hummingbirds, bumblebees, and small bees) and long-tubed populations in the north (mainly small bees), while long-tubed populations in the southwestern range were visited predominantly by hawkmoths (Fig. 2B; Supplemental Table S3). Hawkmoths were the only commonly observed nocturnal pollinator, though a small number of visits (5 total visits) by other moths were recorded at two populations (LVH and SILB12), which constituted only a very small fraction of total visitation. Patterns of pollinator visitation are consistent with those reported by Wenzell et al. (2023), with the exception of hawkmoth visitation to northern population SILB recorded in 2012, which was not observed at SILB during observations conducted in 2013, 2017, or 2018 (in fact, Wenzell et al., 2023 recorded no hawkmoth visitation in any of the 7 studied populations in the northern range over three years of observations).

#### Pollinator exclusion experiment

Results of our day/night pollinator exclusion experiment at nine populations aligned with our expectations that populations with long corollas visited by hawkmoths experience overall higher fruit set at night than during the day, while populations visited by mainly diurnal pollinators show the opposite pattern (Fig. 2C). A GLMM revealed significant differences in fitness for plants exposed to diurnal or nocturnal pollinators, with significant interactions for local pollinators and corolla tube length. For the entire dataset, fitness varied among treatments (χ^2^_2, 420_ = 29.9, p < 0.0001), and treatment interacted with corolla length (χ^2^_2, 420_ = 18.8, p < 0.0001) and with activity period of the most frequent visitor (χ^2^_2, 420_ = 24.4, p < 0.0001). Major pollinator category was also a significant predictor of fruit set independent of treatment (χ^2^_1, 420_ = 4.4, p = 0.035).

Given these significant interaction terms for both corolla length and major pollinator activity period, we proceeded to analyze these groups separately to perform pairwise comparisons of fitness among treatments (Fig. 3). For populations with long corollas visited predominantly by hawkmoths, fitness varied significantly by treatment (χ^2^_2,174_ = 58.059, p < 0.0001). Fitness to plants in the day-open treatment was significantly lower than to those in the night-open (p < 0.0001) and fully open (p < 0.0001) treatments, indicating greater pollination by nocturnal hawkmoth pollinators to long-tubed plants relative to diurnal pollinators (Fig. 3A).

**Figure 3.**
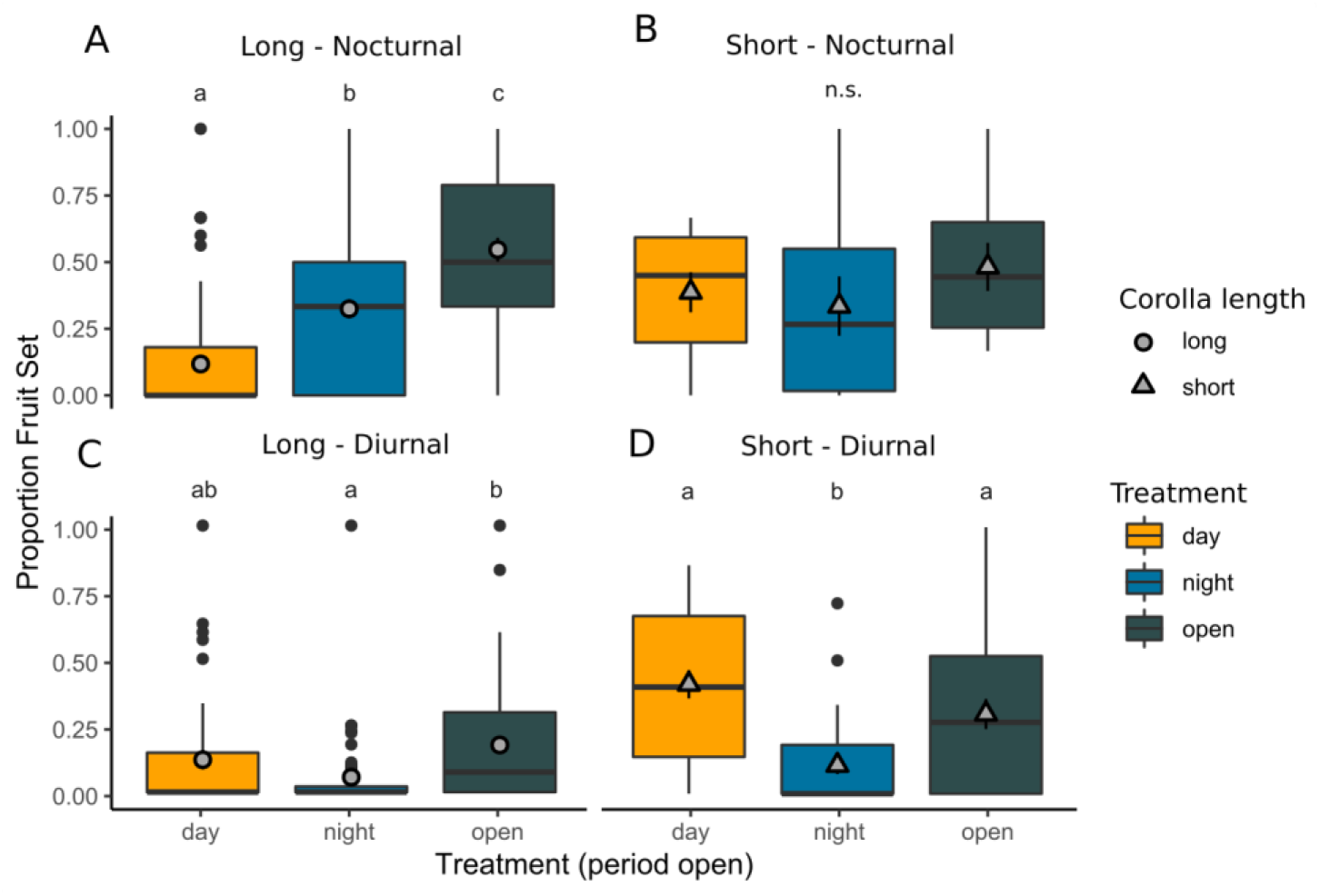
Fitness among exclusion treatments depends on corolla length and major pollinator. Proportion fruit set (fitness) to each pollination treatment for populations grouped by corolla length and activity period of the most frequent visitor: **A**) long corollas, predominately nocturnal visitors; **B**) shorts corollas, predominately nocturnal visitors; **C**) long corollas, predominately diurnal visitors; **D)** short corollas, predominately diurnal visitors. Center bars: median value; upper and lower hinges: first and third quartiles; whiskers: points within 1.5 **×** IQR of hinges; large points: outlying points. Grey points show mean value +/-SE. Treatments with different letters vary significantly in fruit set (p < 0.05) based on GLMM and pairwise tests with Tukey adjustments.

Night-only fitness was also significantly lower than fully open fitness (p = 0.003), suggesting a combination of nocturnal and diurnal pollinators results in higher fruit set overall. In contrast, for short-tubed plants where hawkmoths were the main visitor, no significant differences in fitness were observed among the different pollination treatments (Fig. 3B; χ^2^_2, 26= 1.5, p_ = 0.47), consistent with comparable pollination from both hawkmoths and other diurnal visitors to short-tubed flowers (though we note this combination—short corollas and hawkmoth visitation— occurred at only one population, SIC). For long-tubed populations visited by mostly diurnal visitors, fitness once again varied significantly by treatment (χ^2^_2,136_ = 8.6, p = 0.013), with night-open fitness being significantly lower than fully-open fitness (p = 0.011) and no other significant pairwise differences among treatments (Fig. 3C). For populations with short corollas visited predominantly by diurnal visitors, fitness varied significantly by pollination treatment (χ^2^_2,79_ = 21.9, p < 0.0001), and fitness of plants in the night-open treatment was significantly lower than that of day-open (p < 0.0001) or fully open (p=0.0045) plants (Fig. 3D). Overall, these findings for diurnally-pollinated populations are consistent with expectations that little fruit set was contributed at night where hawkmoths were not observed (though nocturnal pollination was nonetheless above zero in these populations).

#### Populations with long floral tubes experience higher fitness when visited by hawkmoths

Next, we found evidence from a range-wide multi-year dataset that hawkmoth visitation was associated with higher population-level fitness for populations with long corolla tubes but not those with short corolla tubes (Fig. 4A). For long-tubed populations, fitness was greater on average to populations where the most frequent visitor was hawkmoths, compared to those visited by diurnal pollinators (χ^2^_1,161_ = 10.2, p = 0.0014). In contrast, short-tubed populations had no significant difference in population-level fitness whether the most common visitor was nocturnal or diurnal (χ^2^_1,361_ = 0.28, p = 0.6). These findings are consistent with results from our exclusion experiment that hawkmoth pollinators confer increased reproductive fitness only to populations with long corolla tubes.

**Figure 4.**
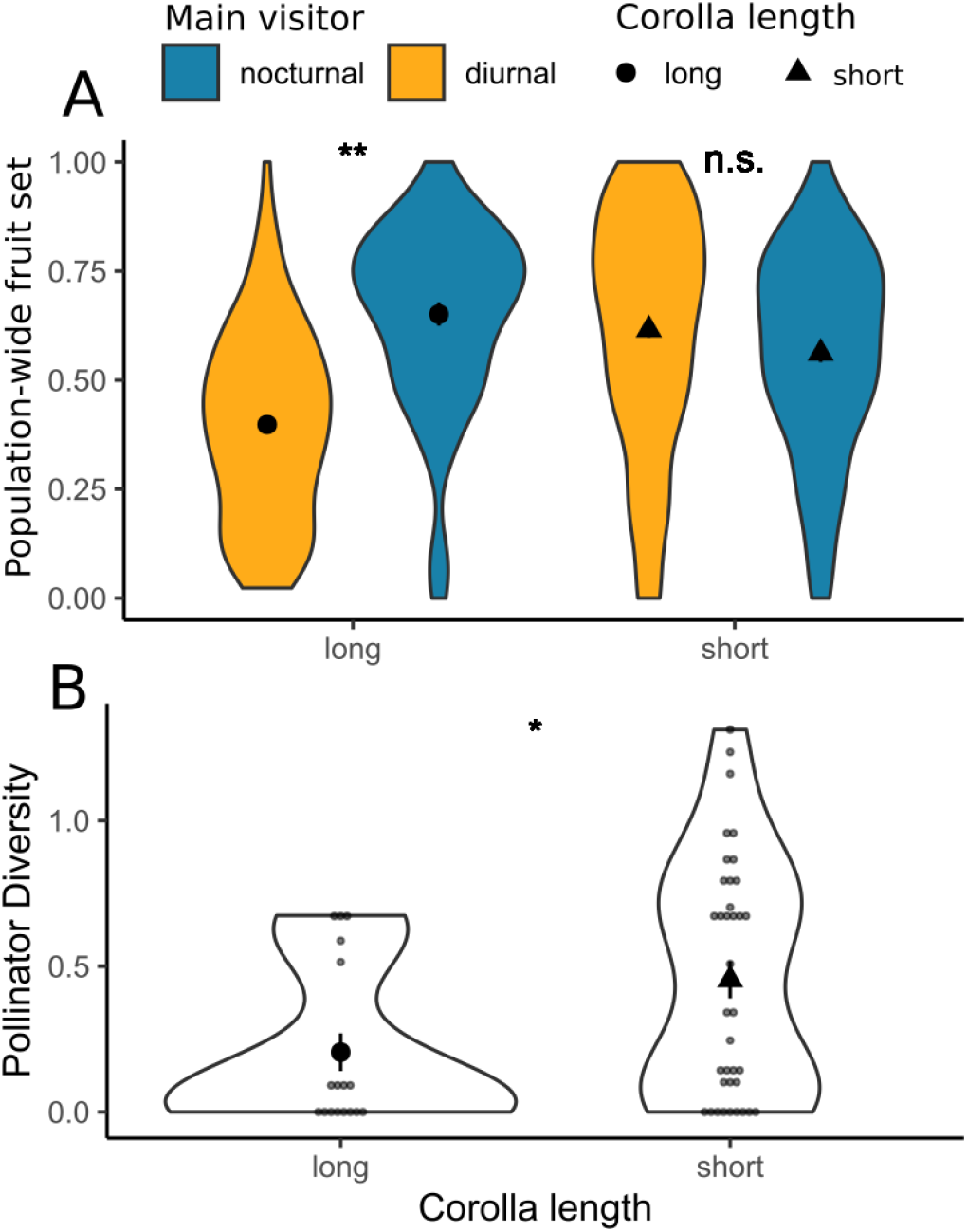
Impacts of corolla length on visitors and fruit set extends to range-wide, multi-year observations in *C. sessiliflora* and the *C. purpurea* complex. **A)** Population-level fitness (proportion fruit set) to populations with long or short corollas whose most frequent floral visitor was either diurnal or nocturnal (i.e., hawkmoths). Asterisks indicates a significant difference in fruit set to populations of the same corolla type when visited by diurnal or nocturnal pollinators (p < 0.05); n.s. indicates no significant difference. **B)** Pollinator diversity (log-transformed Inverse Simpson’s Diversity Index of floral visitors) to populations with either long or short corollas. Asterisk indicates a significant difference among populations by corolla length (p< 0.05). Large central points show mean +/-SE, small points show datapoints.

#### Long corollas act as a filter to limit the diversity of floral visitors

While visitor assemblages to all focal species were generalized overall (Wenzell et al., 2023), we observed that populations of *C. sessiliflora* with long corollas (>41mm on average) typically had visitors from only one or two functional groups, in contrast to more diverse assemblages of most short-tubed populations of both *C. sessiliflora* and the species of the *C. purpurea* complex. Thus, we hypothesized that long floral tubes in *C. sessiliflora* may function as a filter trait, which excludes pollinating taxa without long tongues from accessing nectar rewards. We quantified the diversity of visitors to 23 populations across the ranges of *C. sessiliflora* and the *C. purpurea* complex in relation to their corolla lengths (either long or short), expecting lower levels of visitor diversity to populations with long corollas. In support of this expectation, we found that pollinator diversity was significantly lower in long-tubed populations (Fig. 4B) based on both the GLMM (corolla length: χ^2^_1,50_ = 5.7, p = 0.017; dataset: χ^2^_1,50_ = 0.15, p = 0.7) and Kruskal-Wallis test (KW χ^2^_1,41_ = 6.1, p = 0.013). We did not find evidence that pollinator diversity varied significantly by latitude based on a GLMM (latitude: χ^2^_1,51_ = 2.01, p = 0.16; dataset: χ^2^_1,49= 0.22, p_ = 0.64) and Kruskal-Wallis test (KW χ^2^_22_ = 25.25, p = 0.29; N =41), indicating that lower diversity to long-tubed populations of *C. sessiliflora* is not an artefact of a possible decrease in pollinator diversity due to latitude.

## Discussion

In this study, we present a pollination exclusion experiment across nine populations and analyze a range-wide, multiyear visitation dataset to understand how geographically variable pollinators influence plant reproductive fitness in light of floral trait variation. First, we identify that *C. sessiliflora* is largely self-incompatible and relies on pollinators for reproduction. Next, we find evidence that the distinctive long corollas of *C. sessiliflora* reflect adaptation to hawkmoth pollinators in the southwestern range, where hawkmoths confer increased fitness to plants at night, but only in populations with long corollas. Because this association was not observed in the short-tubed population visited by hawkmoths (SIC), the fitness advantage of hawkmoth pollination appears to depend on long corollas, which we hypothesize increases contact between hawkmoths’ heads and plant sexual organs (Fig. 1C), resulting in higher pollination efficiency, and limits reward access by other potential pollinators. Furthermore, populations visited primarily by diurnal pollinators (including populations of *C. citrina, C. lindheimeri*, the short-tubed, yellow-bracted *C. sessiliflora*, as well as most long-tubed *C. sessiliflora* populations in the north) experienced lower fruit set at night regardless of corolla length, consistent with expectations based on observed floral visitors. Based on our analysis of a previously published range-wide dataset of pollinator visitation to these species (Wenzell et al., 2023), we also found evidence that long-tubed populations have higher reproductive fitness when visited predominantly by hawkmoths compared to diurnal visitors (with no such relationship for short-tubed populations), suggesting long-tubed populations not visited by hawkmoths may face a fitness cost as a result of trait mismatch. Finally, we found that long corolla tubes function as a filter trait to limit the diversity of visitors to *C. sessiliflora*, consistent with a trend toward specialization following adaptation to the most effective pollinator.

Nonetheless, secondary diurnal pollinators (typically pollen-collecting small bees) also contributed significantly to fitness at long-tubed populations across the range, underscoring the role of adaptive generalization in pollinators to provide reproductive assurance in light of geographically variable pollinator assemblages.

The concept of adaptive generalization in floral interactions posits that generalized pollination systems need not necessarily be selected against to allow floral divergence to proceed, but that secondarily important groups of pollinators can also pollinate flowers unless their visitation causes a net fitness negative, at which point theory predicts they would be selected against (Ohashi et al., 2021). During our exclusion experiment, plants with long corollas visited mainly by nocturnal visitors experienced greater fitness when exposed to pollination at night than during the day. This is consistent with hawkmoths (the main nocturnal visitor observed in all populations) being superior pollinators in these populations compared to diurnally active pollinators (Fig. 3A). These findings are aligned with numerous examples in the literature demonstrating associations between increased length of nectar-bearing floral structures (such as corolla tubes or nectar spurs) and pollination by long-tongued pollinators (Anderson and Johnson, 2008), particularly hawkmoths (Fulton and Hodges, 1999; Whittall and Hodges, 2007; Boberg et al., 2014). Nonetheless, fully open treatment plants with long tubes experienced greater fruit set than either night-open or day-open plants, which suggests that the sum of diurnal and nocturnal pollinators contributes greater fitness than either group alone. This finding provides evidence that diurnal secondary pollinators (mainly small bees) provide a significant contribution to fitness and underscores the importance of considering secondary pollinators in studies of floral adaptation (Jaeger et al., 2023). In this system, we propose that small bees represent a secondary pollinator that are not excluded by long corolla tubes (as they collect pollen at the mouths of corolla openings, Wenzell et al., 2023) and that contribute to seed set at a lower level than more effective hawkmoths (when and where they occur) but provide reproductive assurance in portions of the range and in years when hawkmoths are scarce. Thus, we propose this study provides an example of floral divergence via adaptation to the most effective pollinator (hawkmoths in the southwestern range) with important fitness contributions from secondary pollinators (small bees) which provide reproductive assurance through adaptive generalization (Ohashi et al., 2021).

Despite the strong support that hawkmoth visitation confers increased fitness to night-open plants in long-tubed populations, we note that at the long-tubed population SILB12 in 2012, most recorded visitors were in fact hawkmoths, but this did not translate to greater fruit set at night, unlike other long-tubed populations visited by hawkmoths. While this is unexpected, we suspect the overall rarity of hawkmoth visitation in the northern range (Wenzell et al., 2023) and the possibility of pollination by “secondary” diurnal pollinators (likely small bees as at the same population in 2013 (SILB13) and SNRM) contributed to comparable daytime fitness at SILB in 2012. Thus, given the other three long-tubed hawkmoth-visited populations, we feel these results do not invalidate our broader findings that long corolla tubes in *C. sessiliflora* facilitate increased fruit set by hawkmoths in the southwestern range. To this point, we note that in one long-tubed population of *C. sessiliflora* where hawkmoths were the sole recorded visitor in 2019 (SBL), none of the ten plants in the day-open treatment set any fruit whatsoever (e.g., fruit set of 0), and fruit set to the night-open and control treatment plants was nearly identical, suggesting that all contribution to fruit set among treatment plants came exclusively from nocturnal pollinators (Supplemental Figure S2), suggesting that this population is functionally specialized to pollination by hawkmoths (Ollerton et al., 2007).

For the short-tubed populations of *C. sessiliflora*, the pink-flowered population (SIC) was the only population with short tubes whose most frequent visitor was hawkmoths, in addition to small bees and other diurnal visitors such as bee-flies and butterflies (Fig. 2B; Wenzell et al., 2023). Plants at this site experienced no difference in fitness in any pollination treatment, suggesting comparable, effective pollination by both diurnal and nocturnal members of this generalized pollinator assemblage. Given the apparent equal pollination effectiveness of diurnal pollinators at this site, we speculate that this population may represent a scenario where hawkmoths visit and are effective pollinators, but do not exert strong enough selection favoring the evolution of long corollas to overcome potential selection imposed by diverse diurnal pollinators. Put another way, because diurnal pollinators at this site contribute equally to fitness compared to hawkmoths, there is no selection for long corollas to risk filtering out effective diurnal pollinators (particularly bee-flies and butterflies, observed at this site in 2017-2019, which are likely to be excluded from nectar rewards by long corollas, unlike small bees as discussed above). In this case, short corollas are favored to accommodate pollination by generalized diurnal and nocturnal pollinators. This may be in contrast to the scenario favoring long corollas in populations further north and west, where we hypothesize nectar-foraging, diurnal visitors were lacking or ineffective compared to hawkmoths, allowing hawkmoth pollinators to exert selection favoring long corolla tubes (with no net fitness cost, as small bees were not excluded).

In contrast, the other short-tubed population of *C. sessiliflora*, the yellow-flowered SMP, aligned more closely with short-tubed, diurnally-visited populations of *C. citrina* (in floral color and predominant bumblebee visitation) and *C. lindheimeri*, in terms of mainly diurnal visitors and increased fitness conferred during the day, indicating that diurnal pollinators are most effective in these populations. However, we note that nocturnal fruit set was non-zero, suggesting secondary nocturnal pollinators likely contribute here as well, underscoring that nocturnal pollinators are ubiquitous, though often understudied (Hahn and Brühl, 2016). That this short-tubed, yellow-flowered *C. sessiliflora* population aligns with members of the *C. purpurea* complex more closely than with other *C. sessiliflora* populations suggest that it represents a potential pollination ecotype of *C. sessiliflora*, distinct from other conspecific populations. In considering if this scenario could represent potential convergence of yellow flowers in primarily bumblebee-pollinated environments, or rather introgression or common ancestry with the yellow bumblebee-pollinated *C. citrina*, we note that this population is located in the area of overlap between the distributions of *C. sessiliflora* and the yellow-bracted *C. citrina*. Previous genetic analyses did not find evidence that SMP was mis-identified (i.e., in fact a *C. citrina* population) nor that it represents a recent hybrid between *C. sessiliflora* and *C. citrina* (Wenzell et al., 2021). Interestingly, introgression of genes controlling floral pigment leading to adaptive floral color transitions has been reported in *Mimulus* sect. *Diplacus* (Short and Streisfeld, 2023), which could provide one hypothesis explaining how yellow flowers in *C. sessiliflora* may have arisen, especially in areas of overlap with *C. citrina*. Alternatively, ancestral polymorphisms in floral color that precede speciation events (Sánchez-Cabrera et al., 2023) or convergent shifts in floral color among related species (Thomson and Wilson, 2008; Wenzell et al., 2023a; Stone and Wessinger, 2024) provide alternate explanations which require further study.

The observation that long corolla tubes persist in the northern range despite rare hawkmoth visitation may reflect several possibilities: first, evolutionary constraints could limit the ability to reverse elongated floral structures (Huang and Fenster, 2007; Whittall and Hodges, 2007), which may also apply to color (Zufall and Rausher, 2004) as northern *C. sessiliflora* plants lack floral pigmentation compared to southern populations. Second, the current pollinator community of the northern range may not reflect the conditions under which this species evolved, as recent land use change and habitat fragmentation may be more dramatic in the tall grass prairies of *C. sessiliflora*’s northern range (strongly impacted by intensive agriculture) than the rangelands it inhabits elsewhere (Templeton et al., 2001). This could also impact local pollinator populations, potentially driving declines in populations of hawkmoths or other nocturnal pollinators. Another possibility is the history of glaciation which impacted the northern portion of the range to a greater extent than the southern—potentially allowing the populations of the southern range extent more time to diversify and evolve to local ecological conditions (Dalton et al., 2020). Additionally, it is possible that long corolla tubes could be selected for and/or maintained due to selection against herbivores/ seed predators (Strauss and Whittall, 2006), which could be hypothesized to oviposit more successfully on short-tubed flowers compared to long, another possibility which warrants further study.

Finally, we acknowledge several limitations to this study. Because pollinator observations did not extend throughout the night but ended shortly after last light, pollinators other than the observed hawkmoths may have visited night-open plants and contributed to fruit set, and therefore we cannot attribute all fruit set in night-open plants exclusively to pollination by hawkmoths. Additionally, we acknowledge the day-versus-night timing of exclusion treatments is an imperfect proxy to gauge fruit set contributed by specific pollinator groups, as hawkmoths are known to sometimes forage during the day (Stöckl and Kelber, 2019), and diurnal visitors such as hummingbirds and bees may forage at dawn and dusk, potentially outside of the “day-open” treatment window. Nonetheless, we found overwhelmingly that relative fitness among treatments matched expectations based on observed visitors at the population-level, which reinforces that our observations largely reflected visitation to treatment plants. Overall, our results support the hypothesis that when they are visited by hawkmoths, long-tubed populations of *C. sessiliflora* experience greater fitness during periods when hawkmoths are known to be active than at other times. These findings are consistent with the notion that long corolla tubes in *C. sessiliflora* are associated with increased fitness to plants when visited by hawkmoths, and thus may represent an adaptation to pollination by hawkmoths, a pollination mode which has not previously been demonstrated in the genus *Castilleja*.

## Conclusions

Overall, this study highlights the continuum of pollination modes and levels of generalization and specialization that can exist within a species across wide geographic distributions. We present evidence that nocturnal hawkmoths are superior pollinators to long-tubed populations of *C. sessiliflora*, that long-tubed populations visited predominantly by hawkmoths have higher fitness, and that long tubes limit the diversity of floral visitors, resulting in a potential fitness cost due to trait mismatch in parts of the range where hawkmoths are scarce. While some populations showed conditions that could facilitate evolution toward specialization (i.e.,, SBL, where hawkmoths were the only effective pollinator), others remained generalized (e.g., SIC, where hawkmoths and diurnal visitors contributed equally to fitness), and yet others are at risk of trait mismatch with local pollinators (the northern SILB and SNRM). This study underscores that small bees represent important secondary pollinators capable of pollinating flowers regardless of tube length, which likely provide reproductive assurance and may not be selected against, consistent with adaptive generalization in pollination systems. Thus, by comparing pollination by different pollinator groups across the range of a widespread species and its parapatric congeners, considering natural variation in a key floral trait, this study provides evidence that trends toward specialization in pollination can emerge from generalist pollinated environments, but nonetheless are still characterized by important secondary pollinators and potential trait mismatch due to geographic variation in local pollinators.

## Acknowledgements

The authors thank A. Iler and H. Briggs for input on the study and analyses and D. Tank and M. Egger for insight on the study systems. We are grateful for field assistance from C. Woolridge, A.J. Morgan, M. Rhodes, E. Hilpman, and S. Todd. We thank the following permitting agencies for permission and access to study sites: The Nature Conservancy Chapters of TX and IL, Native Prairies Association of Texas, City of Lubbock, TX, TX Parks and Wildlife Department, NM Energy, Minerals and Natural Resources Department, IL Nature Preserves Commission, IL Department of Natural Resources, and US Forest Service. Funding was provided by the American Philosophical Society Lewis and Clark Grant, Friends of Nachusa Grasslands, Northwestern University Program in Plant Biology and Conservation, and an NSF Graduate Research Fellowship award (DGE-1842165 to K.E.W.). Additional support was provided by the Negaunee Institute for Plant Conservation and Action at the Chicago Botanic Garden. The authors declare no conflict of interest.

## Author contributions

K.E.W., J.B.F., and K.A.S. conceived of the study; K.E.W., J.B.F., J.Y.Z., and K.A.S. collected and interpreted the data; K.E.W. performed statistical analyses and drafted the manuscript with input from J.B.F., J.Y.Z., and K.A.S.

## LITERATURE CITED

Anderson, B., and S. D. Johnson. 2008. The geographical mosaic of coevolution in a plant-pollinator mutualism. Evolution; International Journal of Organic Evolution 62: 220–225.

Barrett, S. C. H. 2013. The evolution of plant reproductive systems: how often are transitions irreversible? Proceedings of the Royal Society B: Biological Sciences 280.

Boberg, E., R. Alexandersson, M. Jonsson, J. Maad, J. Ågren, and L. A. Nilsson. 2014. Pollinator shifts and the evolution of spur length in the moth-pollinated orchid Platanthera bifolia. Annals of Botany 113: 267–275.

Campbell, D. R., N. M. Waser, and E. J. Melendez-Ackerman. 1997. Analyzing Pollinator-Mediated Selection in a Plant Hybrid Zone: Hummingbird Visitation Patterns on Three Spatial Scales. The American Naturalist 149: 295–315.

Chuang, T. I., and L. R. Heckard. 1991. Generic Realignment and Synopsis of Subtribe Castillejinae (Scrophulariaceae-Tribe Pediculareae). Systematic Botany 16: 644–666.

Crepet, W. L., K. C. Nixon, and M. A. Gandolfo. 2004. Fossil evidence and phylogeny: the age of major angiosperm clades based on mesofossil and macrofossil evidence from Cretaceous deposits. American Journal of Botany 91: 1666–1682.

Dalton, A. S., M. Margold, C. R. Stokes, L. Tarasov, A. S. Dyke, R. S. Adams, S. Allard, et al. 2020. An updated radiocarbon-based ice margin chronology for the last deglaciation of the North American Ice Sheet Complex. Quaternary Science Reviews 234: 106223.

Faegri, K., and L. van der Pijl. 1971. Principles of Pollination Ecology. Elsevier.

Fenster, C. B., W. S. Armbruster, P. Wilson, M. R. Dudash, and J. D. Thomson. 2004. Pollination Syndromes and Floral Specialization. Annual Review of Ecology, Evolution & Systematics 35: 375–403.

Fox, J., S. Weisberg, B. Price, D. Adler, D. Bates, G. Baud-Bovy, B. Bolker, et al. 2020. car: Companion to Applied Regression.

Fulton, M., and S. A. Hodges. 1999. Floral isolation between Aquilegia formosa and Aquilegia pubescens. Proceedings of the Royal Society of London B: Biological Sciences 266: 2247–2252.

Hahn, M., and C. A. Brühl. 2016. The secret pollinators: an overview of moth pollination with a focus on Europe and North America. Arthropod-Plant Interactions 10: 21–28.

Hartig, F. 2016. DHARMa: residual diagnostics for hierarchical (multi-level/mixed) regression models. Website https://CRAN.R-project.org/package=DHARMa [accessed 22 May 2022].

Ho, W. W., and S. D. Smith. 2016. Molecular evolution of anthocyanin pigmentation genes following losses of flower color. BMC Evolutionary Biology 16: 98.

Huang, S., and C. B. Fenster. 2007. Absence of Long-Proboscid Pollinators for Long-Corolla-Tubed Himalayan Pedicularis Species: Implications for the Evolution of Corolla Length. International Journal of Plant Sciences 168: 325–331.

Jaeger, S., C. Girvin, N. Demarest, and E. LoPresti. 2023. Secondary pollinators contribute to reproductive success of a pink-flowered sand verbena population. Ecology 104: e3977.

Johnson, S. D. 2006. Pollinator-driven speciation in plants. In L. D. Harder, and S. C. H. Barrett [eds.], Ecology and Evolution of Flowers, 295–310. Oxford University Press.

Kay, K. M., and R. D. Sargent. 2009. The Role of Animal Pollination in Plant Speciation: Integrating Ecology, Geography, and Genetics. Annual Review of Ecology, Evolution, and Systematics 40: 637–656.

Lázaro, A., R. Lundgren, and Ø. Totland. 2009. Co-flowering neighbors influence the diversity and identity of pollinator groups visiting plant species. Oikos 118: 691–702.

Lenth, R. 2018. lsmeans: Least-Squares Means.

Magnusson, A., H. Skaug, A. Nielsen, C. Berg, K. Kristensen, M. Maechler, K. van Bentham, et al. 2020. glmmTMB: Generalized Linear Mixed Models using Template Model Builder.

Miller, R. B. 1981. Hawkmoths and the Geographic Patterns of Floral Variation in Aquilegia caerulea. Evolution 35: 763–774.

Miller, T. J., R. A. Raguso, and K. M. Kay. 2014. Novel adaptation to hawkmoth pollinators in Clarkia reduces efficiency, not attraction of diurnal visitors. Annals of Botany 113: 317–329.

Nesom, G. L., and M. Egger. 2014. Review of the Castilleja purpurea complex (Orobanchaceae). Phytoneuron 15: 1–16.

van der Niet, T., and S. D. Johnson. 2012. Phylogenetic evidence for pollinator-driven diversification of angiosperms. Trends in Ecology & Evolution 27: 353–361.

Ohashi, K., A. Jürgens, and J. D. Thomson. 2021. Trade-off mitigation: a conceptual framework for understanding floral adaptation in multispecies interactions. Biological Reviews 96: 2258–2280.

Ollerton, J., A. Killick, E. Lamborn, S. Watts, and M. Whiston. 2007. Multiple meanings and modes: on the many ways to be a generalist flower. TAXON 56: 717–728.

R Development Core Team. 2021. R: a language and environment for statistical computing: reference index. R Foundation for Statistical Computing, Vienna.

Rausher, M. D. 2008. Evolutionary Transitions in Floral Color. International Journal of Plant Sciences 169: 7–21.

Sánchez-Cabrera, M., E. Narbona, M. Arista, P. L. Ortiz, F. J. Jiménez-López, A. Fuller, B. Carter, and J. B. Whittall. 2023. Can a flower color ancestral polymorphism transcend speciation? Evolutionary Biology.

Sargent, R. D., and S. P. Otto. 2006. The role of local species abundance in the evolution of pollinator attraction in flowering plants. The American Naturalist 167: 67–80.

Short, A. W., and M. A. Streisfeld. 2023. Ancient hybridization leads to the repeated evolution of red flowers across a monkeyflower radiation. Evolution Letters: qrad024.

Stebbins, G. L. 1970. Adaptive Radiation of Reproductive Characteristics in Angiosperms, I: Pollination Mechanisms. Annual Review of Ecology and Systematics 1: 307–326.

Stöckl, A. L., and A. Kelber. 2019. Fuelling on the wing: sensory ecology of hawkmoth foraging. Journal of Comparative Physiology A 205: 399–413.

Stone, B. W., and C. A. Wessinger. 2024. Ecological Diversification in an Adaptive Radiation of Plants: The Role of De Novo Mutation and Introgression. Molecular Biology and Evolution 41: msae007.

Strauss, S., and J. B. Whittall. 2006. Non-pollinator agents of selection on floral traits. In L. D. Harder, and S. C. H. Barrett [eds.], Ecology and Evolution of Flowers, 120–138. Oxford University Press.

Tank, D. C., J. M. Egger, and R. G. Olmstead. 2009. Phylogenetic Classification of Subtribe Castillejinae (Orobanchaceae). Systematic Botany 34: 182–197.

Templeton, A. R., R. J. Robertson, J. Brisson, and J. Strasburg. 2001. Disrupting evolutionary processes: The effect of habitat fragmentation on collared lizards in the Missouri Ozarks. Proceedings of the National Academy of Sciences 98: 5426–5432.

Thompson, J. N. 2005. The Geographic Mosaic of Coevolution. University of Chicago Press.

Thomson, J. D., and P. Wilson. 2008. Explaining Evolutionary Shifts between Bee and Hummingbird Pollination: Convergence, Divergence, and Directionality. International Journal of Plant Sciences 169: 23–38.

Tripp, E. A., and P. S. Manos. 2008. Is Floral Specialization an Evolutionary Dead-End? Pollination System Transitions in Ruellia (acanthaceae). Evolution 62: 1712–1737.

Wang, B., Z.-Y. Tong, Y.-Z. Xiong, X.-F. Wang, W. S. Armbruster, and S.-Q. Huang. 2023. The evolution of flower-pollinator trait matching, and why do some alpine gingers appear to be mismatched? Annals of Botany: mcad141.

Waser, N. M., L. Chittka, M. V. Price, N. M. Williams, and J. Ollerton. 1996. Generalization in Pollination Systems, and Why it Matters. Ecology 77: 1043–1060.

Wei, N., R. L. Kaczorowski, G. Arceo-Gómez, E. M. O’Neill, R. A. Hayes, and T.-L. Ashman. 2021. Pollinators contribute to the maintenance of flowering plant diversity. Nature: 1–5.

Wenzell, K. E., A. J. McDonnell, N. J. Wickett, J. B. Fant, and K. A. Skogen. 2021. Incomplete reproductive isolation and low genetic differentiation despite floral divergence across varying geographic scales in Castilleja. American Journal of Botany 108: 1270–1288.

Wenzell, K. E., M. Neequaye, and K. J. R. P. Byers. 2023a. Do color polymorphisms reflect pollinator shifts?: Integrating floral traits as signals to a novel pollinator to understand species divergence. 2023.10.29.564637.

Wenzell, K. E., K. A. Skogen, and J. B. Fant. 2023b. Range-wide floral trait variation reflects shifts in pollinator assemblages, consistent with pollinator-mediated divergence despite generalized visitation. Oikos 2023: e09708.

Whittall, J. B., and S. A. Hodges. 2007. Pollinator shifts drive increasingly long nectar spurs in columbine flowers. Nature 447: 706–709.

Zufall, R. A., and M. D. Rausher. 2004. Genetic changes associated with floral adaptation restrict future evolutionary potential. Nature 428: 847–850.

