## SupplementalMaterials for "Adaptive generalization in pollinators: hawkmoths increase fitness to long-tubed flowers, but secondary pollinators remain important"

### SUPPLEMENT

#### Breeding system of *Castilleja sessiliflora*

**Table S1.** Fruit set to plants growing in a common garden that were either fully bagged to exclude pollinators and unmanipulated ("bagged" flowers) or bagged and hand-pollinated with self-pollen ("hand-selfed" flowers). Plants were grown at Chicago Botanic Garden from seed wild-collected from populations across the range. Tests were conducted in 2019.

| Population | State | Exclusion experiment population? | No. of plants | No. of flowers bagged | No. of fruits to bagged flowers | Bagged fruit set (percent) | No. of flowers hand self-pollinated | No. of fruits to hand-selfed flowers | Hand-selfed fruit set (percent) |
|---|---|---|---|---|---|---|---|---|---|
| SMP | TX | Yes | 1 | 29 | 0 | 0.00% | 12 | 0 | 0% |
| SCC | CO | Yes | 1 | 9 | 0 | 0.00% | 3 | 0 | 0% |
| SDC | CO | No | 2 | 19 | 0 | 0.00% | 8 | 0 | 0% |
| SILB | IL | Yes | 1 | 35 | 1 | 2.86% | 6 | 0 | 0% |
| SFP | MN | No | 1 | 13 | 0 | 0.00% | 4 | 0 | 0% |
| **Total** | | | **6** | **105** | **1** | **0.95%** | **33** | **0** | **0%** |

**Table S2.** Fruit set to plants growing in natural populations that were fully bagged to exclude pollinators during pollinator exclusion experiments conducted in 2012 and 2013.

| Population | State | Year | No. of plants | No. of flowers bagged | No. of fruits to bagged flowers | Bagged fruit set (percent) |
|---|---|---|---|---|---|---|
| SCC | CO | 2012 | 24 | 91 | 3 | 3.30% |
| SDC | CO | 2012 | 24 | 69 | 0 | 0.00% |
| SILB | IL | 2012 | 6 | 26 | 0 | 0.00% |
| SILB | IL | 2013 | 28 | 311 | 0 | 0.00% |
| **Total** | | | **82** | **497** | **3** | **0.60%** |

**Table S3**. Pollinator visitation data for focal populations by count of visits, proportion of total visits, and visitation rate (number of visits/ flower/ hour).

**Table S3. Pollinator visitation to exclusion experiment populations**

| | | | Count | | | | | | |
|---|---|---|---|---|---|---|---|---|---|
| Population | Year | Dataset | Other diurnal | Hawkmoth | Small/ med bees | Other nocturnal | Bumble-bee | Humming-bird | Total visits |
| SCC | 2012 | Narrow view | 1 | 88 | 30 | 0 | 0 | 0 | 119 |
| SILB12 | 2012 | Narrow view | 0 | 34 | 0 | 3 | 0 | 0 | 37 |
| SILB13 | 2013 | Narrow view | 0 | 0 | 3 | 0 | 0 | 0 | 3 |
| CQL | 2019 | Narrow view | 0 | 0 | 1 | 0 | 0 | 0 | 1 |
| CQL | 2019 | Wide view | 0 | 0 | 7 | 0 | 67 | 29 | 103 |
| LVH | 2019 | Narrow view | 0 | 0 | 1 | 0 | 0 | 3 | 4 |
| LVH | 2019 | Wide view | 5 | 3 | 1 | 2 | 0 | 44 | 55 |
| SBL | 2019 | Narrow view | 0 | 4 | 0 | 0 | 0 | 0 | 4 |
| SBL | 2019 | Wide view | 0 | 29 | 0 | 0 | 0 | 0 | 29 |
| SCL | 2019 | Narrow view | 0 | 46 | 2 | 0 | 0 | 0 | 48 |
| SCL | 2019 | Wide view | 0 | 16 | 0 | 0 | 2 | 0 | 18 |
| SIC | 2019 | Narrow view | 8 | 53 | 20 | 0 | 0 | 0 | 81 |
| SIC | 2019 | Wide view | 3 | 49 | 0 | 0 | 0 | 0 | 52 |
| SMP | 2019 | Narrow view | 0 | 1 | 0 | 0 | 0 | 0 | 1 |
| SMP | 2019 | Wide view | 0 | 0 | 0 | 0 | 105 | 0 | 105 |
| SNRM | 2019 | Narrow view | 1 | 0 | 165 | 0 | 62 | 0 | 228 |
| SNRM | 2019 | Wide view | 0 | 0 | 7 | 0 | 14 | 0 | 21 |

| | | | Proportion | | | | | |
|---|---|---|---|---|---|---|---|---|
| Population | Year | Dataset | Other diurnal | Hawkmoth | Small/ medium bees | Other nocturnal | Bumble-bee | Humming-bird |
| SCC | 2012 | Narrow view | 0.008403 | 0.739496 | 0.252101 | 0 | 0 | 0 |
| SILB12 | 2012 | Narrow view | 0 | 0.918919 | 0 | 0.081081 | 0 | 0 |
| SILB13 | 2013 | Narrow view | 0 | 0 | 1 | 0 | 0 | 0 |
| CQL | 2019 | Narrow view | 0 | 0 | 1 | 0 | 0 | 0 |
| CQL | 2019 | Wide view | 0 | 0 | 0.067961 | 0 | 0.650485 | 0.2815534 |
| LVH | 2019 | Narrow view | 0 | 0 | 0.25 | 0 | 0 | 0.75 |
| LVH | 2019 | Wide view | 0.090909 | 0.054545 | 0.018182 | 0.036364 | 0 | 0.8 |
| SBL | 2019 | Narrow view | 0 | 1 | 0 | 0 | 0 | 0 |
| SBL | 2019 | Wide view | 0 | 1 | 0 | 0 | 0 | 0 |
| SCL | 2019 | Narrow view | 0 | 0.958333 | 0.041667 | 0 | 0 | 0 |
| SCL | 2019 | Wide view | 0 | 0.888889 | 0 | 0 | 0.111111 | 0 |
| SIC | 2019 | Narrow view | 0.098765 | 0.654321 | 0.246914 | 0 | 0 | 0 |
| SIC | 2019 | Wide view | 0.057692 | 0.942308 | 0 | 0 | 0 | 0 |
| SMP | 2019 | Narrow view | 0 | 1 | 0 | 0 | 0 | 0 |
| SMP | 2019 | Wide view | 0 | 0 | 0 | 0 | 1 | 0 |
| SNRM | 2019 | Narrow view | 0.004386 | 0 | 0.723684 | 0 | 0.27193 | 0 |
| SNRM | 2019 | Wide view | 0 | 0 | 0.333333 | 0 | 0.666667 | 0 |

| | | | Visitation rate (visits/ flower/ hour) | | | | | |
|---|---|---|---|---|---|---|---|---|
| Population | Year | Dataset | Other diurnal | Hawkmoth | Small/ medium bees | Other nocturnal | Bumble-bee | Humming-bird |
| SCC | 2012 | Narrow view | 0.001484 | 0.130628 | 0.044532 | 0 | 0 | 0 |
| SILB12 | 2012 | Narrow view | 0 | 0.04629 | 0 | 0.004084 | 0 | 0 |
| SILB13 | 2013 | Narrow view | 0 | 0 | 0.001938 | 0 | 0 | 0 |
| CQL | 2019 | Narrow view | 0 | 0 | 0.000995 | 0 | 0 | 0 |
| CQL | 2019 | Wide view | 0 | 0 | 0.000665 | 0 | 0.006361 | 0.0027531 |
| LVH | 2019 | Narrow view | 0 | 0 | 0.001948 | 0 | 0 | 0.0058442 |
| LVH | 2019 | Wide view | 0.001513 | 0.000908 | 0.000303 | 0.000605 | 0 | 0.0133148 |
| SBL | 2019 | Narrow view | 0 | 0.007084 | 0 | 0 | 0 | 0 |
| SBL | 2019 | Wide view | 0 | 0.060421 | 0 | 0 | 0 | 0 |
| SCL | 2019 | Narrow view | 0 | 0.085928 | 0.003736 | 0 | 0 | 0 |
| SCL | 2019 | Wide view | 0 | 0.0156 | 0 | 0 | 0.00195 | 0 |
| SIC | 2019 | Narrow view | 0.011483 | 0.076077 | 0.028708 | 0 | 0 | 0 |
| SIC | 2019 | Wide view | 0.000979 | 0.015985 | 0 | 0 | 0 | 0 |
| SMP | 2019 | Narrow view | 0 | 0.000408 | 0 | 0 | 0 | 0 |
| SMP | 2019 | Wide view | 0 | 0 | 0 | 0 | 0.005737 | 0 |
| SNRM | 2019 | Narrow view | 0.001082 | 0 | 0.178571 | 0 | 0.0671 | 0 |
| SNRM | 2019 | Wide view | 0 | 0 | 0.007799 | 0 | 0.015597 | 0 |

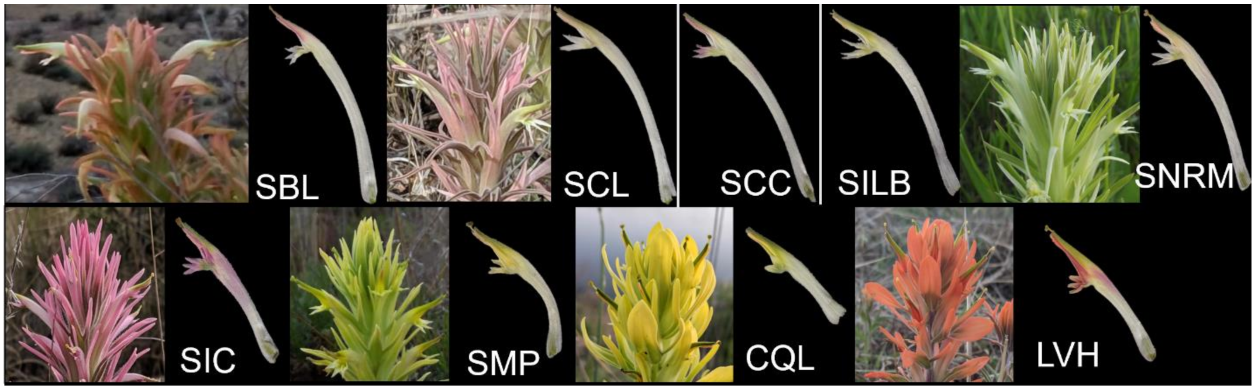


**Figure S1.** Representative photos of floral morphs and corollas for populations where exclusion experiments were conducted. Top row: long-tubed populations; bottom row: short-tubed populations. First letter of population codes denotes species: S = *C. sessiliflora*, C = *C. citrina*, L = *C. lindheimeri*. Images not to scale.

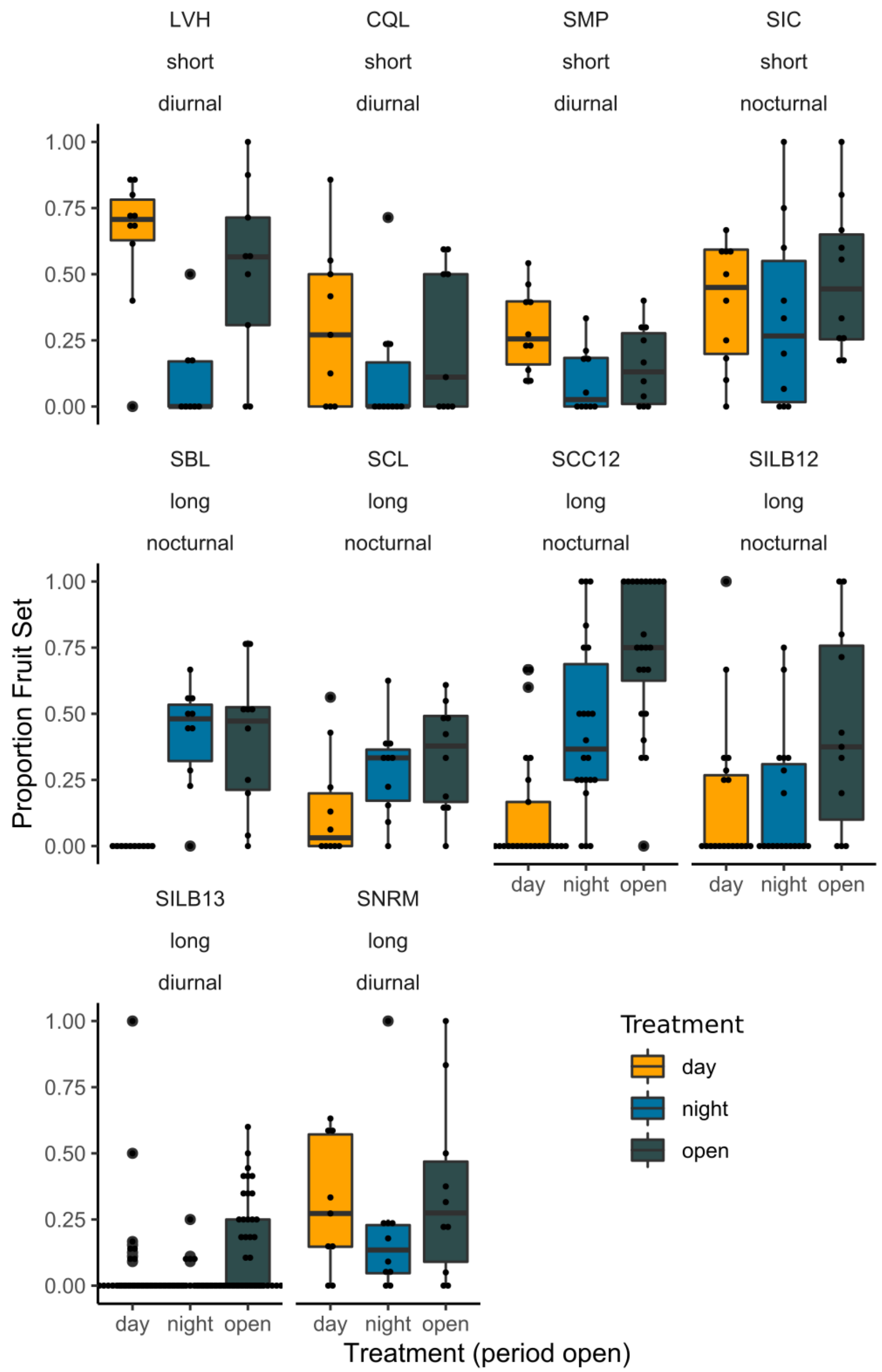


**Figure S2.** Population-level results of the pollinator exclusion experiment performed in nine natural populations. Proportion fruit set is shown for each treatment: plants open to pollinators during the day, at night, or fully open. Population codes are given at the top of each plot. Plots are labelled with the corolla length morph of the population and whether the most frequent visitor to the population during experimental windows was diurnal or nocturnal (i.e., hawkmoths). The exclusion experiment was repeated in subsequent years at SILB (2012 and 2013).
